# Human alveolar Type 2 epithelium transdifferentiates into metaplastic KRT5+ basal cells during alveolar repair

**DOI:** 10.1101/2020.06.06.136713

**Authors:** Jaymin J. Kathiriya, Chaoqun Wang, Alexis Brumwell, Monica Cassandras, Claude Le Saux, Paul Wolters, Michael Matthay, Harold A Chapman, Tien Peng

**Affiliations:** Department of Medicine, University of California San Francisco

**Keywords:** Human stem cells, alveolar type 2 cells (AEC2s), metaplasia, idiopathic pulmonary fibrosis (IPF), KRT5+, basal cells, alveolar-basal intermediate cell (ABI)

## Abstract

Understanding differential lineage potential of orthologous stem cells across species can shed light on human disease. Here, utilizing 3D organoids, single cell RNAseq, and xenotransplants, we demonstrate that human alveolar type 2 cells (hAEC2s), unlike murine AEC2s, are multipotent and able to transdifferentiate into KRT5+ basal cells when co-cultured with primary fibroblasts in 3D organoids. Trajectory analyses and immunophenotyping of epithelial progenitors in idiopathic pulmonary fibrosis (IPF) indicate that hAEC2s transdifferentiate into metaplastic basal cells through alveolar-basal intermediate (ABI) cells that we also identify in hAEC2-derived organoids. Modulating hAEC2-intrinsic and niche factors dysregulated in IPF can attenuate metaplastic basal cell transdifferentiation and preserve hAEC2 identity. Finally, hAEC2s transplanted into fibrotic immune-deficient murine lungs engraft as either hAEC2s or differentiated KRT5+ basal cells. Our study indicates that hAEC2s-loss and expansion of alveolar metaplastic basal cells in IPF are causally connected, which would not have been revealed utilizing murine AEC2s as a model.

**Highlights:**

- Human AEC2s transdifferentiate into KRT5+ basal cells when accompanied by primary adult human lung mesenchyme in 3D organoid culture.
- Alterations of hAEC2-intrinsic and niche factors dysregulated in IPF can modify metaplastic hAEC2 transdifferentiation.
- hAEC2s engraft into fibrotic lungs of immune-deficient mice and transdifferentiate into metaplastic basal cells.
- Transcriptional trajectory analysis suggests that hAEC2s in IPF gives rise to metaplastic basal cells via alveolar-basal intermediate cells.

## Introduction

The use of murine tissue-resident stem cells to model human stem cell behavior relies on the assumption that lineage capacity is conserved across species. Understanding differences in the behavior of orthologous stem cell populations across species could shed light on human diseases where a particular stem cell is implicated. Murine genetic studies have shown that mouse alveolar type 2 cells (mAEC2s) are the resident stem cell population in the alveoli that constitute the entire gas exchange surface of the lung (Barkauskas et al., 2013; Rock et al., 2011). In idiopathic pulmonary fibrosis (IPF), the most deadly and prevalent form of diffuse parenchymal lung disease, hAEC2s are lost from the alveoli and replaced with metaplastic KRT5+ basal cells that normally appear in the conducting airways (Adams et al., 2019; Habermann et al., 2019; Smirnova et al., 2016; Xu et al., 2016). Rigorous murine genetic lineage tracing has shown that metaplastic KRT5+ cells in the alveoli are not derived from mAEC2s, but rather from KRT5-/SOX2+ progenitors in the mouse airway after severe injury (Kumar et al., 2011; Ray et al., 2016; Xi et al., 2017; Yang et al., 2018). However, it is not clear whether a similar population in the human airway exists that contributes to metaplastic basal cells, as the airways contain key anatomic differences across the two species. The question of where metaplastic KRT5+ basal cells come from in human IPF has not been answered, but it is clinically significant because the presence of alveolar KRT5+ basal cells directly correlates with mortality in IPF (Prasse et al., 2019). In this study, we made a surprising finding that hAEC2s, but not mAEC2s, can readily transdifferentiate into KRT5+ basal cells in organoid culture and xenotransplant. These results implicate hAEC2s as the source of metaplastic KRT5+ basal cells in IPF, and our 3D organoid and xenotransplant models provide powerful platforms to study human epithelial metaplasia in IPF that is not possible with murine stem cells.

## RESULTS

### Primary adult human lung mesenchyme drives human AEC2 transdifferentiation into KRT5+ basal cells *in vitro*

Previously reported hAEC2 organoids have utilized MRC5, a fetal human lung fibroblasts cell line, as feeders to maintain hAEC2s *in vitro* (Barkauskas et al., 2013; Li et al., 2020; Zacharias et al., 2018). We initially hypothesized that co-culture with primary adult human lung mesenchyme (AHLM) would improve the growth of hAEC2 organoids compared to MRC5. To test this, we isolated AEC2s from normal human donor lungs (age greater than 18, no underlying lung disease at time of organ donation) and co-cultured them with MRC5 or AHLM isolated from the same donor lung in our heterotypic 3D organoid system. We utilized the HTII-280 antibody to isolate hAEC2s (EpCAM+/HTII-280+) (Fig. S1A), and a separate flow cytometry strategy to isolate AHLM (EpCAM-/CD45-/CD11b-/CD31-) that was previously described by scRNAseq as predominantly PDGFRa+ fibroblasts (Wang et al., 2018). To confirm the purity of the hAEC2 isolation by anti-HTII-280, we performed cytospin and scRNAseq of the EpCAM+/HTII-280+ population to confirm a homogenous population expressing common hAEC2 markers such as *SFTPC, LAMP3, ABCA3, and NAPSA*, and near absence of cells with basal, club, goblet, and ciliated cell markers (Fig. 1A and S1B). As previously shown, co-culture of hAEC2s with MRC5 resulted in numerous SFTPC+ organoids (Fig. 1A, B, and S1C). But much to our surprise, we saw a dramatic loss of SFTPC accompanied by gradual appearance of KRT5, an airway basal cell marker, in hAEC2 organoids co-cultured with AHLM at Day 7 (D7) and 14 of co-culture (Fig. 1A and B, and S1C). By D14 of co-culture, the majority cells in organoids derived from hAEC2-AHLM co-culture were KRT5+, whereas the majority of cells in the hAEC2-MRC5 co-cultured remained SFTPC+. Furthermore, isolation of epithelial cells from the 3D organoid culture containing AHLM vs. MRC5 showed a significant reduction of *SFTPC* expression concurrent with a dramatic increase in *KRT5* expression in the organoids co-cultured with AHLM (Fig. 1C). A parallel experiment using mAEC2s cocultured with primary adult murine lung mesenchyme (AMLM) produced no KRT5+ organoids (Fig.1A and B, and S1C).

**Figure 1.**
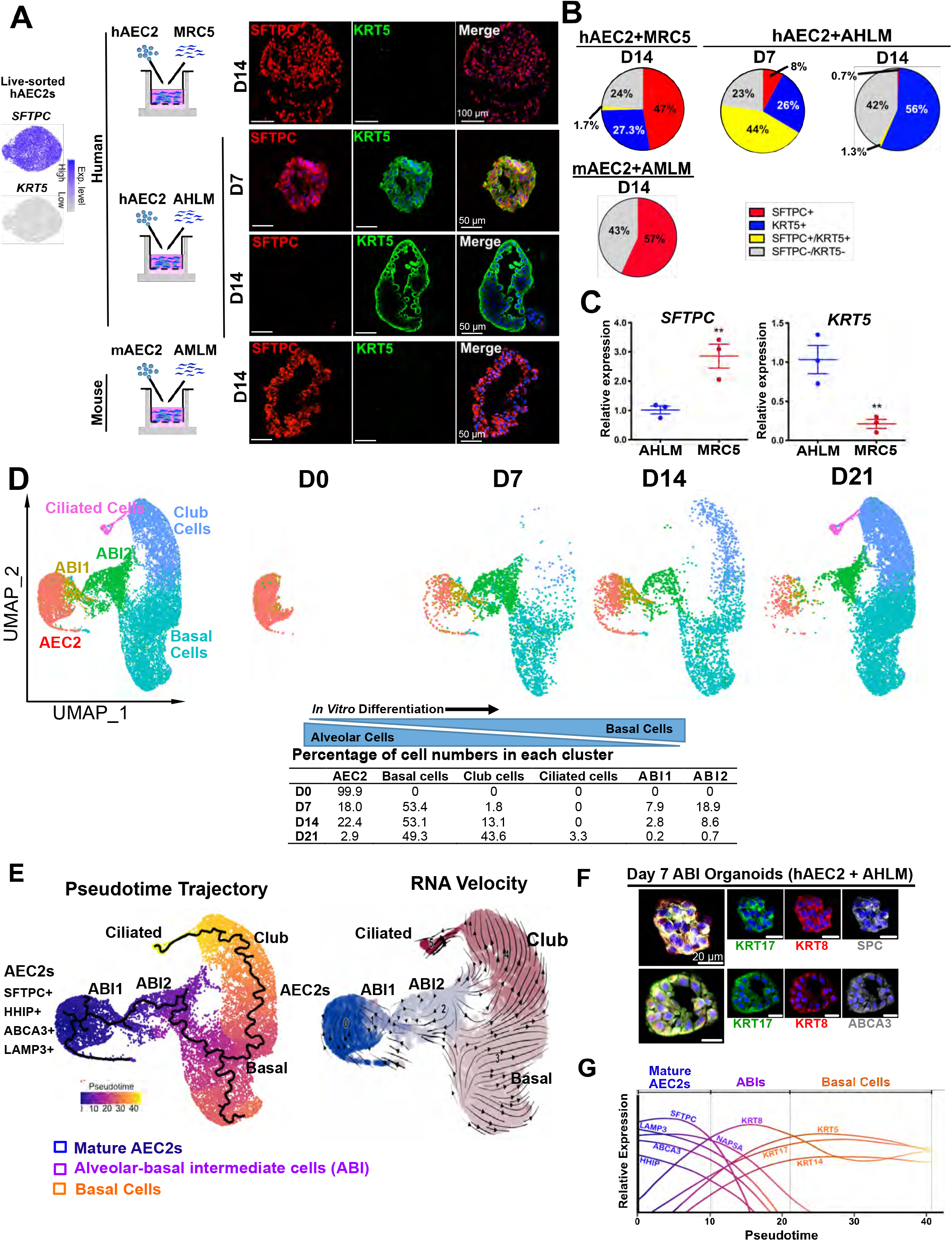
Primary adult human lung mesenchyme drives human AEC2 transdifferentiation into KRT5+ basal cells *in vitro*. (A) Live/EpCAM+/HTII-280+ cells (hAEC2s) were sorted and sequenced, confirming purified *SFTPC+* population. These cells were mixed with either primary adult human lung mesenchyme (AHLM) or MRC5 fetal human lung fibroblast cell line in 1:6 epithelial:mesenchyme ratio. Mouse AEC2s (mAEC2s) were sorted GFP+ cells from the lung of adult *Sftpc^CreERT2/+^:R26^RmTmG/+^* mouse and co-cultured with the adult mouse lung mesenchyme (AMLM) taken from the same lung. (B) Quantification of cell composition in the organoids. (C) mRNA analysis of hAEC2s organoids co-cultured with AHLM vs. MRC5 fibroblasts. (D) scRNA-seq of freshly sorted HTII-280+ hAEC2s (D0) and hAEC2s co-cultured with AHLM at D7, D14, and D21. (E) Pseudotime ordering of epithelial cells from D0, D7, D14, and D21 organoids identify an hAEC2s → ABIs → Basal → Club → Ciliated cell trajectory based on marker analysis over pseudotime trajectory that was confirmed on RNA Velocity. (F) Staining of D7 hAEC2+AHLM organoids identify presence of KRT17+/KRT8+/SFTPC+ and KRT17+/KRT8+/ABCA3+ ABIs. (G) Summary plot of pseudotime gene expression levels as cell transition from mature hAEC2s → ABIs → Basal cell states. Abbr.: hAEC2 (human alveolar type 2 cells), AHLM (adult human lung mesenchyme), ABI (alveolar-basal intermediate) See also Figure S1 and S2.

Histological analysis of organoids co-cultured with AHLM at various time points revealed presence of SFTPC+/KRT5+ dual positive cells and mature alveolar or basal cells at D7 and D14 in addition to a fraction of cells that stained for neither (Fig. 1A and B, and S1C). We then performed a time-course scRNAseq of organoids at D7, D14, and D21. Epithelial cells from D0 (live sort of HTII-280+ prior to culture, Fig. 1A and S1A, B), D7, D14, and D21 organoids were merged and clustered by uniform manifold approximation and projection (UMAP) where 6 main clusters emerged (Fig. 1D and S1D). Four clusters were easily identified by lineage markers of hAEC2s, basal cells, club cells, and ciliated cells respectively. Two clusters showed intermediate lineage phenotypes with both AEC2 and basal cell markers, indicating the presence of an alveolar-basal intermediate cell (ABI) population. Time course shows a time-dependent loss of the hAEC2 population and gain of KRT5+ basal cell population, along with an early emergence of ABIs at D7 that eventually recedes with time in culture (Fig. 1D and S1D). This indicates that hAEC2 transdifferentiates into basal cells through intermediate cellular states with both hAEC2 and basal cell traits. Concurrent with the disappearance of the ABIs with time, there is also a progressive emergence of SCGB1A1+ club cells and FOXJ1+ ciliated population in culture, showing that transdifferentiated KRT5+ cells are functional basal cells undergoing further differentiation into club and ciliated lineages with time.

We then performed Monocle pseudotime trajectory analysis of the various epithelial clusters arising in the scRNAseq, which shows a dominant trajectory arising from hAEC2s to basal cells through an ABI population, characterized by the step-wise loss of alveolar markers and accompanied by gradual acquisition of basal markers (Fig. 1E,G, S1E, S2A). Visualization of gene expression along the hAEC2 transdifferentiation trajectory shows that well-established AEC2 marker, *LAMP3*, and novel hAEC2 marker, *HHIP*, are quickly lost first as mature hAEC2s transition into ABIs. The emergence of early basal marker, *KRT17*, and intermediate marker, *KRT8* (Jiang et al., 2020; Kobayashi et al., 2019; Strunz et al., 2019), concurrent with the persistence of AEC2 markers *SFTPC, ABCA3*, and *NAPSA*, marks the ABI population most prominently seen in early culture at D7 (Fig. S1E, S2A). The presence of the ABIs is confirmed on immunophenotyping of D7 organoids, where we see KRT17+/KRT8+/SFTPC+ and KRT17+/KRT8+/ABCA3+ cells (Fig. 1F), which are not present in the normal human lung (Fig. S1F). As transdifferentiation progresses, the ABIs give away to mature basal cells as all hAEC2 markers are lost with the emergence of mature basal cell markers *KRT5* and *KRT14* in D14 organoids (Fig. S2A,B). The pseudotime analysis and RNA Velocity analysis (Fig. 1E) also shows a clear trajectory from the hAEC2-derived basal cells to SCGB1A1+ club cells and then FOXJ1+ ciliated cells, the presence of which is further confirmed by histology (Fig. S2C). These findings indicate that hAEC2s possess remarkable plasticity compared to mAEC2s, with the capacity to transdifferentiate and generate a diverse repertoire of airway epithelium that is influenced by mesenchymal niche factors.

### hAEC2-intrinsic and niche factors altered in IPF modulate metaplastic AEC2 transdifferentiation

Ingenuity Pathway Analysis (IPA) on the single cell trancriptomes of hAEC2s cocultured with AHLM showed that BMP activation is downregulated in hAEC2s in culture, concurrent with the upregulation of airway/basal cell differentiation pathways regulated by SOX2 and TP63 (Fig. 2A). We and others have previously shown that secreted BMP antagonists are upregulated in the mesenchyme of IPF lungs (Adams et al., 2019; Cassandras et al., 2019; Koli et al., 2006; Myllarniemi et al., 2008) (Fig. S3A), where ectopic basal cells appear in the alveoli accompanied by the loss of hAEC2s. To test the hypothesis that BMP antagonists are dysregulated in cultured AHLM when compared to MRC5, we first examined the transcriptome of AHLM vs. MRC5 by bulk RNAseq. Differential gene expression analysis shows significant upregulation of multiple secreted BMP antagonists in the AHLM, whereas BMP ligands were preferentially upregulated in MRC5 (Fig. 2B). Addition of recombinant BMP4 to hAEC2 organoids co-cultured with AHLM significantly attenuated *KRT5* expression while increasing *SFTPC* expression in the organoids (Fig. 2C), which is confirmed by immunophenotyping of organoid sections (Fig. 2D). This demonstrates that BMP activation in the hAEC2 niche maintains AEC2 fate, and upregulation of BMP antagonism could promote KRT5+ metaplasia seen in IPF.

**Figure 2.**
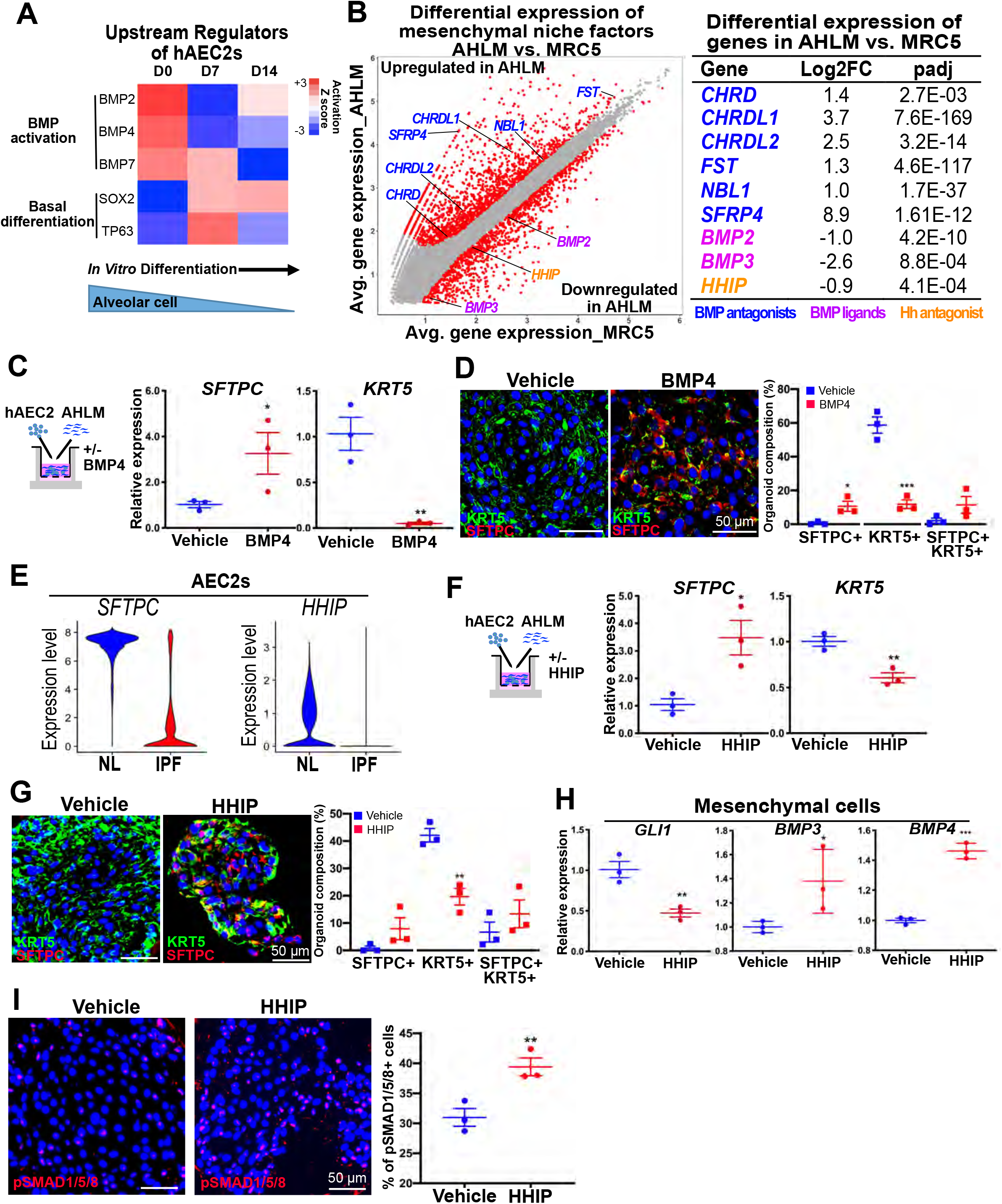
hAEC2-intrinsic and niche factors altered in IPF modulate metaplastic hAEC2 transdifferentiation. (A) IPA upstream regulator analysis of hAEC2s co-cultured with AHLM. (B) Bulk RNA-seq analysis of MRC5 and AHLM with differential gene expression analysis. (C) Treatment of hAEC2s+AHLM organoids with BMP4 increases *SFTPC* and decreases *KRT5* mRNA expression at D14 of culture. (D) BMP treatment increases SFTPC+ cells and decreases KRT5+ cells in D14 organoids. (E) hAEC2s from IPF lungs demonstrates decreased *SFTPC* and *HHIP* expression when compared to hAEC2s from normal donors. (F) Treatment of hAEC2s+AHLM organoids with HHIP increases *SFTPC* and decreases *KRT5* mRNA expression at D14 of culture. (G) HHIP increases SFTPC+ cells and decreases KRT5+ cells in D14 organoids. (H) HHIP-treated AHLM isolated from D14 organoid culture have decreased *GLI1* and increased *BMP3* and *BMP4* mRNA levels. (I) HHIP-treated hAEC2s+AHLM organoids at D14 show significantly higher fraction of pSMAD1/5/8+ cells. See also Figure S3

Differential gene expression analysis demonstrated that *HHIP* was downregulated in AHLM *in vitro* (Fig. 2B). *HHIP* encodes for hedgehog-interacting protein, a decoy receptor of hedgehog (Hh) ligand that attenuates Hh activation by sequestering free Hh ligands (Chuang et al., 2003). As demonstrated in the organoid scRNAseq, *HHIP* is highly expressed in mature hAEC2 (Fig. 1E and S1E and 3B), contrasting with the expression pattern of its murine orthologue, *Hhip*, which is predominantly expressed in the mesenchyme with little expression in mAEC2s (Fig. S3B). To assess the expression of *HHIP* in IPF, we performed scRNAseq of the epithelium isolated from patients undergoing lung transplantation for IPF along with healthy control donors. Analysis of the cluster sizes demonstrates a dramatic loss of hAEC2s and gain of KRT5+ basal cells in IPF (Fig. S3C), which is confirmed by cytospin of the EpCAM+ cells (Fig. S3E). The hAEC2s remaining in IPF express a significantly reduced quantity of *HHIP* compared to hAEC2s from controls (Fig. 2E), which we validated in a larger dataset of single cell analysis of IPF and controls (Adams et al., 2019) (Fig. S3A). Addition of recombinant HHIP to hAEC2 organoids co-cultured with AHLM significantly attenuated *KRT5* expression while increasing *SFTPC* expression in the organoids (Fig. 2F), which is confirmed by immunophenotyping of organoid sections (Fig. 2G). HHIP treatment attenuated Hh activation in the hAEC2 niche, concurrent with an increase in the expression of BMP ligands (Fig. 2H). HHIP-treated organoids also demonstrated a significant increase of phospho-SMAD1/5/8+ cells (Fig. 2I), a readout for BMP activation. IPA upstream regulator analysis suggests that BMP activation is downregulated in IPF hAEC2s (Fig. S3D). These results demonstrate that a Hh-BMP signaling loop altered in IPF could modify metaplastic AEC2 transdifferentiation *in vitro*.

**Figure 3.**
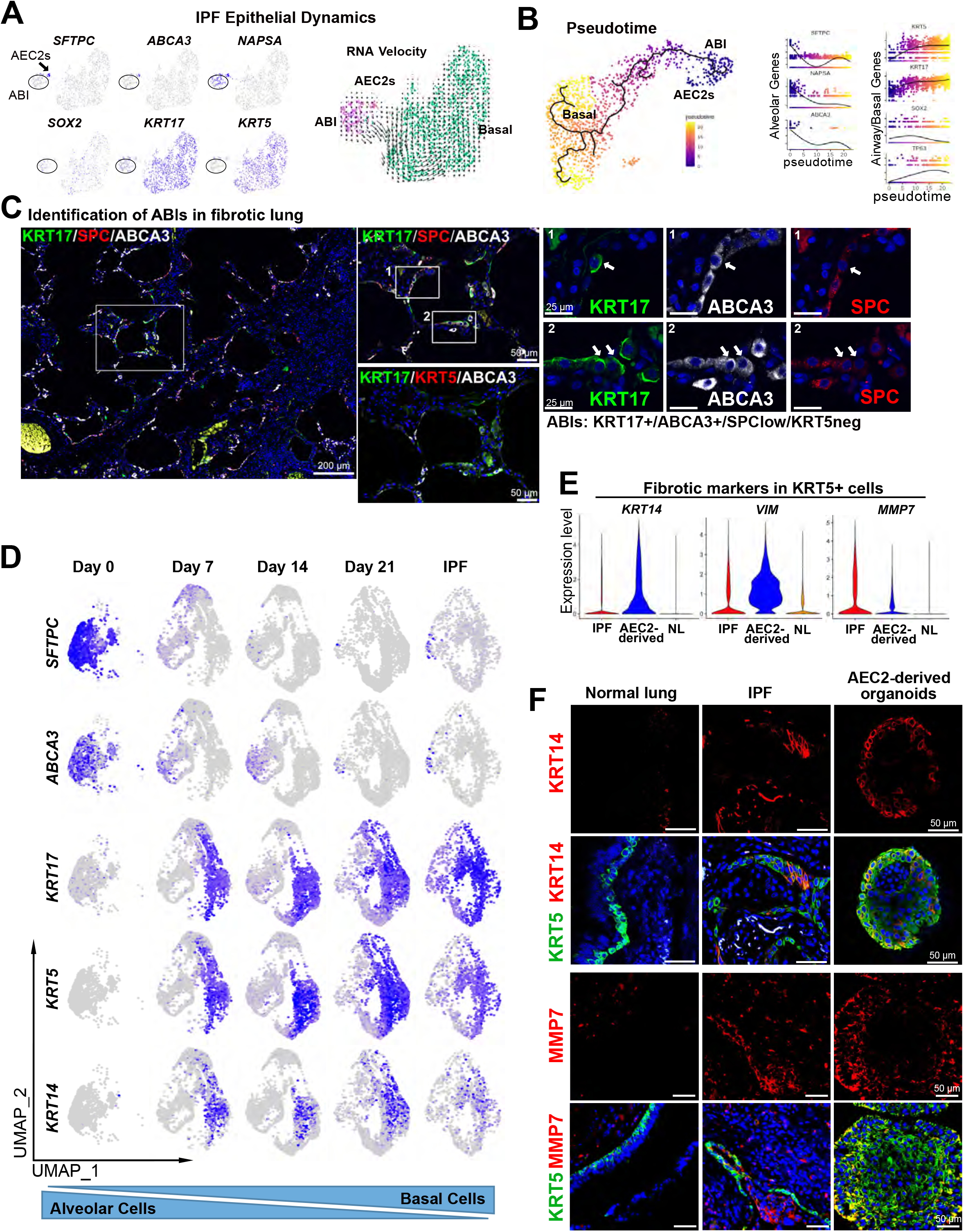
Evidence of hAEC2 transdifferentiation into basal cells through alveolar-basal intermediate cells in IPF. (A) scRNA-seq of IPF EpCAM+ cells shows a small SFTPC+ hAEC2 population in addition to SFTPC^low^/ABCA3^low^/KRT17+ ABI population, along with an expanded KRT17+/KRT5+ basal cells. RNA velocity analysis shows a predicted lineage from hAEC2s to mature KRT17+/KRT5+ basal cells through an ABI population. (B) Pseudotime ordering of IPF epithelial cells shows a similar differentiation from mature hAEC2s → ABIs → Basal cell population. (C) Staining of IPF lung identify KRT17+/ABCA3+/SFTPC^low^ ABIs in the region between heavily fibrotic and normal looking alveolar regions of the lung. Serial sections were stained for KRT5, showing these cells are KRT5^neg^. The image is a composite image of multiple images taken at 20X and stitched together (See STAR Methods). (D) Comparison of scRNA-seq analysis of hAEC2-derived organoids at different time points along with IPF lungs. (E) Violinplots show increased expression of KRT14, Vimentin, and MMP7 in both IPF and hAEC2-derived organoid basal cells but not in normal lung basal cells. (F) Staining of IPF lung and hAEC2-derived organoids identified presence of KRT14+ and MMP7+ basal cells (KRT5+) unlike normal lung. See also Figure S4.

### Evidence of hAEC2 transdifferentiation into basal cells through alveolar-basal intermediates (ABIs) in IPF

In IPF, hAEC2s are gradually lost and replaced with metaplastic KRT5+ basal cells (Fig S3C,E) in the alveoli (Xu et al., 2016), which correlates with poor prognosis. To determine the potential lineage relationship between the epithelial progenitors in injured human lungs, we performed various trajectory analyses to infer potential lineage relationships amongst single cell subsets of EpCAM+ cells from IPF lungs. Majority of the EpCAM+ cells in IPF lungs are *KRT5+* and *KRT17+*. The residual *SFTPC+* cells coclustered with other known AEC2 genes including *ABCA3*, and *NAPSA* (Fig. 3A) in addition to *SOX*2 and *KRT17* (Fig. 3A) with minimal expression of *KRT5*, identifying an ABI population similar to the ones identified in the organoids. RNA velocity demonstrated a transcriptional trajectory from hAEC2s towards the *KRT5+* basal population through a population of ABIs (Fig. 3A). Monocle pseudotime analysis demonstrates a clear path between hAEC2s and *KRT5+* basal cells that goes through a *SFTPC^low^/ABCA3^low^/KRT17+/KRT5*-ABI population as previously seen in our organoid analysis (Fig. 3B and S4B). Over pseudotime, the alveolar genes are progressively lost and airway genes including basal cell markers *KRT17* and *KRT5* appear (Fig 3B), consistent with our organoid data. Immunophenotyping of IPF lungs shows the presence of ABCA3+/SFTPC^low^ cells within alveolar-like and micro-honeycombed areas (Fig. 3C). These cells were also KRT17+ but KRT5-, consistent with the ABI transcriptome we observed in organoid (Fig. 1D,E). Such cells are not found in normal lungs (Fig. S1F). These immunophenotyping results are consistent with two recent studies compiling cellular transcriptome atlases of IPF lungs that identified transitional cell types resembling hAEC2s and basal cells (termed KRT5-/KRT17+ or aberrant basaloid cells by the two studies respectively) specifically in the IPF lungs (Adams et al., 2019; Habermann et al., 2019).

We observe that overall AEC2-derived epithelial populations in organoids gradually shifts towards a pathogenic IPF-like epithelium, as judged by decreased AEC2-specific markers and increased basal cell markers (Fig. 3D). This analysis suggested a similarity in epithelial make up of hAEC2-derived organoids and IPF lungs. To compare basal cells in IPF, normal lungs, and hAEC2-derived organoids, we segregated out *KRT5+* cells from these distinct datasets and re-clustered them by UMAP (Fig. S4C). To identify common features of IPF and hAEC2-derived basal cells in comparison to normal lung basal cells, we compared the transcriptomes of IPF basal cells and hAEC2-derived basal cells with normal lung basal cells, generating a list of differentially expressed genes that were shared between IPF and hAEC2-derived basal cells (Fig 3E and S4D). Further analysis of these genes identified several IPF-associated genes including *MMP1, MMP7, KRT14*, and *VIM* (Fig. S4D)(Rosas et al., 2008; Smirnova et al., 2016; Xu et al., 2016). Consistent with this analysis, we also identified presence of *MMP7, KRT14*, and *VIM* in both parenchymal basal cells in IPF lungs and in hAEC2-derived organoids, but not in normal lungs (Fig. 3F and S4E). These results implicate deregulated hAEC2s as a source of metaplastic basal cells observed in the parenchyma of IPF lungs.

### hAEC2 is capable of metaplastic KRT5+ transdifferentiation in a fibrotic host *in vivo*

To determine whether hAEC2s are capable of KRT5+ differentiation *in vivo*, we transplanted them into the fibrotic lungs of immune-deficient murine hosts. We first injured NSG mice deficient for B-cell, T-cell, and NK cell, with bleomycin at Day 0 (D0), followed at day 10 by oropharyngeal injection of sorted EpCAM+/HTII-280+ hAEC2s isolated from normal donor lungs. We then analyzed the lungs for engraftment of human cells at D20 (Fig. 4A). Using the human-specific nuclear antigen (HNA), we were able to identify patches of engrafted human cells predominantly in the damaged alveolar regions. Co-staining with KRT5 identified HNA+/KRT5+ areas where the metaplastic basal cells reconstitute cystic-like structures seen in IPF lungs (Fig. 4B). Utilizing an antibody specific to human-pro-SP-C, we also note presence of HNA+/pro-SP-C+ hAEC2 patches in the lung that are spatially distinct from dysplastic basal patches (Fig. 4B). Immunohistology analysis showed that the majority of the engrafted hAEC2s patches retained alveolar differentiation (~75%), with a minority undergoing metaplastic basal cell differentiation (~25%) (Fig. 4C). Next, we repeated the transplantation of hAEC2s alongside AHLM to determine the metaplastic differentiation of engrafted hAEC2s. We transplanted the hAEC2:AHLM at 2:1 ratio. Co-transplant of AHLM significantly increased the ratio of engrafted human patches undergoing metaplastic basal cell differentiation, with a majority of the patches constituting KRT5+ basal cells (~80%) and a minority retaining alveolar differentiation (20%) (Fig. 4C). Furthermore, we also identified the presence of rare KRT17+/KRT8+/SPC^low^ ABIs within otherwise SPC+ patches (Fig. 4D). Within the HNA+/KRT5+ patches, we observed hAEC2-derived HNA+/SCGB1A1+ club cells and HNA+/Acetylated Tubulin 1A+ ciliated cells, indicating differentiation of hAEC2s into multiple airway lineages (Fig. 4E). Finally, consistent with the pathologic nature of hAEC2-derived organoids and IPF basal cells, hAEC2-derived basal cells in xenografts also stained for KRT14, MMP7, and VIM (Fig. 4F, Fig. S4F). The xenotransplantation experiments demonstrate that hAEC2s are capable of reconstituting a fibrotic milieu *in vivo* and transdifferentiate into KRT5+ basal cells, presenting a novel humanized model of epithelial metaplasia in lung fibrosis (Fig. 4G).

**Figure 4.**
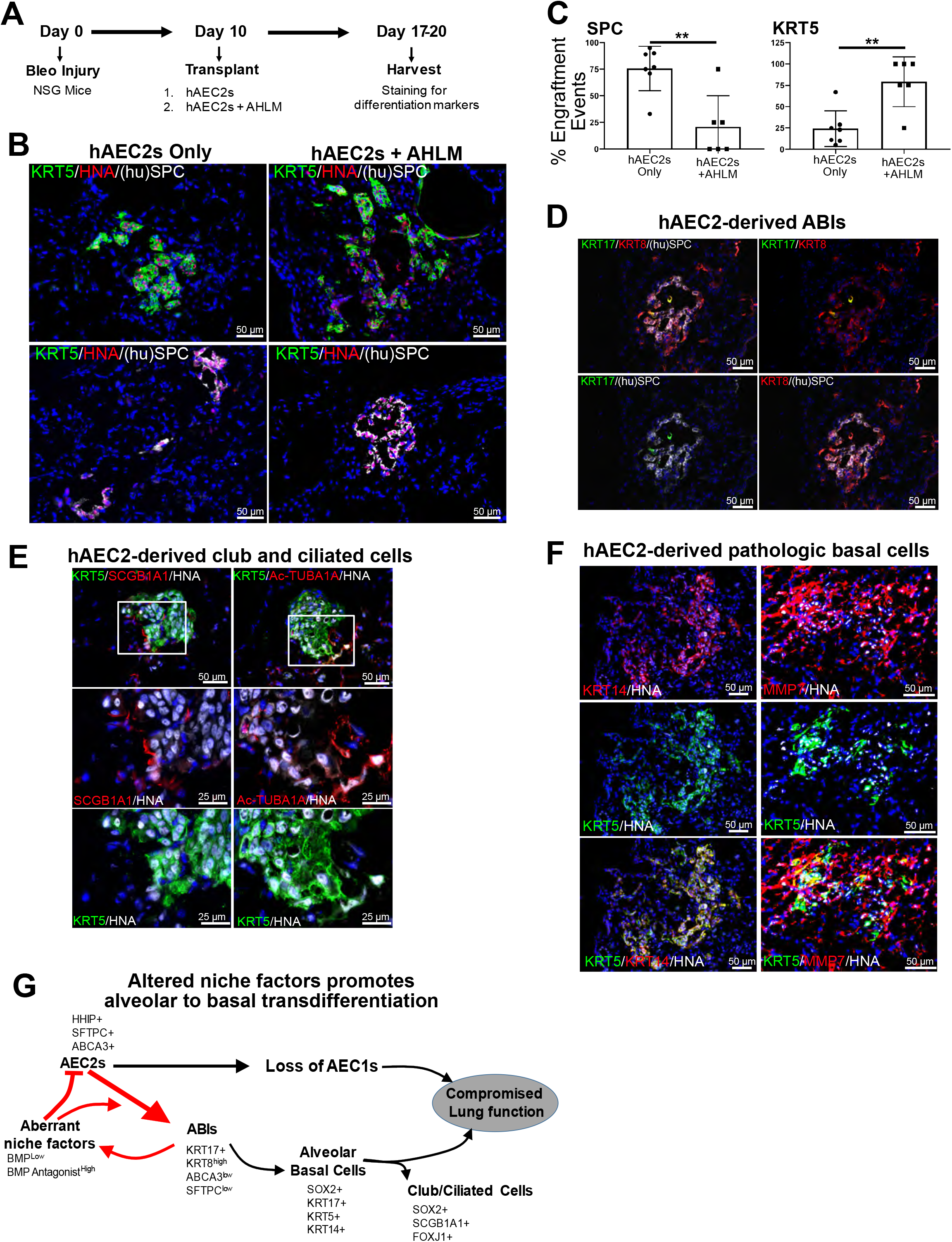
hAEC2 is capable of metaplastic KRT5+ transdifferentiation in a fibrotic host *in vivo*. (A) Experimental setup of xenotransplant experiment testing engraftment of hAEC2s. (B) hAEC2s engrafted in the injured lungs and differentiated towards either KRT5+ (green) basal cells or pro-SFTPC+ (white) AEC2s. (C) Co-transplantation mesenchyme reduced the number of AEC2 patches observed in the lungs of mice while increasing basal cell differentiation. Each dot represents the number of engrafted regions (> 5 cells) in one section with 3 mice analyzed per condition. (D) Co-staining of KRT17, KRT8, and SFTPC of xenografts identify rare KRT17+/KRT8+/SFTPC^low^ ABIs at D10 post transplantation. (E) SCGB1A1+ club cells (red) or Ac-TubA1A+ multiciliated cells (red) are found near hAEC2-derived basal cells. (F) hAEC2s-derived basal cells in xenografted mouse lungs are KRT14+ and MMP7+. (G) Altered mesenchymal niche promotes aberrant epithelial cell dynamics causing alveolar to basal cell differentiation in IPF. See also Figure S4.

## DISCUSSION

While it is well known that the total number of hAEC2s in lungs of IPF patients decreases with disease progression and metaplastic basal cells expand (Habermann et al., 2019; Smirnova et al., 2016; Xu et al., 2016), it has been unclear whether and how these marked changes in epithelial dynamics in IPF are mechanistically linked. The unexpected finding that hAEC2s robustly transdifferentiate into functional basal cells with cues from “activated” mesenchymal cells invited the possibility that a parallel process occurs in IPF lungs, possibly accounting for the known loss of hAEC2s and metaplastic basal cell expansion in IPF. On a single cell level, there is a clear trajectory that appears incremental from mature hAEC2 to KRT5+ basal cells (Fig. 3A, B). Consistent with these progressions is the finding of hybrid ABIs expressing both alveolar and airway proteins in tissues of IPF. We identified one such intermediate, KRT17+/ABCA3+/SPClow cells, as a discrete, early reprogramming intermediate both in the epithelial transcriptional analyses and *in situ* within areas of microhoneycombing in IPF lungs (Fig. 3C). Another group of recent studies have also identified an intermediate cell-state between AEC2 to AEC1 differentiation that is marked by elevated expression of KRT8 and increased TGFb1 signaling, which persists in IPF lungs (Jiang et al., 2020; Kobayashi et al., 2019; Strunz et al., 2019). Interestingly, we find similar features in ABIs, suggesting that this intermediate state may not be unique only during AEC2-to-AEC1 transdifferentiation but also during AEC2-to-Basal transdifferentiation and that upregulation of KRT8 and TGFb1 signaling may represent signaling events contributing to metaplastic hAEC2 differentiation. Collectively, these findings support the view that AEC2s in IPF are subverted from their normal role as self-renewing progenitors and a source of AEC1s, which likely compromises normal alveolar stability and contributes to progressive loss of lung function (Fig. 4G).

A critical element in the diversion of hAEC2s from their role in maintenance of normal alveolar epithelial lining is altered input from their mesenchymal niche. A comparative analysis of the RNA transcriptomes of MRC5 and AHLM reveals extensive differences in pathways known to affect hAEC2 differentiation between the two mesenchymal cell types (Fig. 2B). Unlike MRC5 cells, AHLM cells placed in 3D organoids upregulate a group of secretory products that antagonize BMP signaling and downregulate HHIP, an inhibitor of sonic hedgehog (SHH) signaling. BMP signaling is known to promote murine AEC2 differentiation whereas SHH signaling favors differentiation of murine epithelial progenitors to KRT5+ basal cells by promoting BMP antagonism (Cassandras et al., 2019; Wang et al., 2018). The addition of HHIP or BMP4 to the 3D organoid co-cultures attenuates airway differentiation and maintains hAEC2 lineage, consistent with a causal impact of altered BMP and SHH signaling on hAEC2 fate.

Human AEC2s are capable of either expanding to replenish alveolar epithelial cells or reprogramming to airway-like basal cells and their known mature progeny, as confirmed *in vivo* by transplantation of uncultured, purified hAEC2s into immunodeficient, bleomycin-injured mice (Fig. 4B). However, it is unclear under what situations this multipotency is an advantage for hAEC2 function. In the short term, the acquisition of basal-like features by AEC2s likely conveys increased mobility and proliferative potential that could facilitate maintenance of crucial epithelial barriers in the face of AEC death. If so, this potential appears to be a two-edge sword because continued diversion of hAEC2s into an airway phenotype likely contributes to the accumulation of micro-honeycombed structures lined by airway cells in alveolar regions prominent in fibrotic lung diseases and associated with disease progression. It remains to be determined if and under what circumstances AEC2 reprogramming toward metaplastic basal cells in the alveoli in the setting of disease is reversible. The organoid and transplant platforms used here to recreate conditions supporting either airway or alveolar differentiation of hAEC2s *in vivo* should be a powerful approach to develop modulating factors such as BMPs that preserve AEC2 fate, or HHIP that redirect mesenchymal cells to support AEC2 expansion and possibly improve lung function.

## Author Contributions

J.K., C.W., H.C., and T.P. conceived the experiments and wrote the manuscript. C.W., J.K., A.B., C.L.S., M.C., M.M., and P.W. performed the experiments, collected samples, and analyzed data.

## Acknowledgements

We thank Alisha Baldwin for providing technical assistance; Parnassus Flow Cytometry Core for assistance with cell sorting for bulk and single cell RNA analysis (P30DK063720); Eunice Wan and the Institute for Human Genetics Core for processing of single cell RNA samples and high-throughput sequencing. GEO accession number for raw RNA sequencing data is listed in Materials and Methods. This work is supported by NIH grants DP2AG056034, R01HL142552, and Tobacco Related Disease Research Program New Investigator Award T.P., R01HL128484 and U01HL134766 to H.C., and Nina Ireland Program Award to M.M. for human lung collection.

---

**Figure S1 related to Figure 1:**
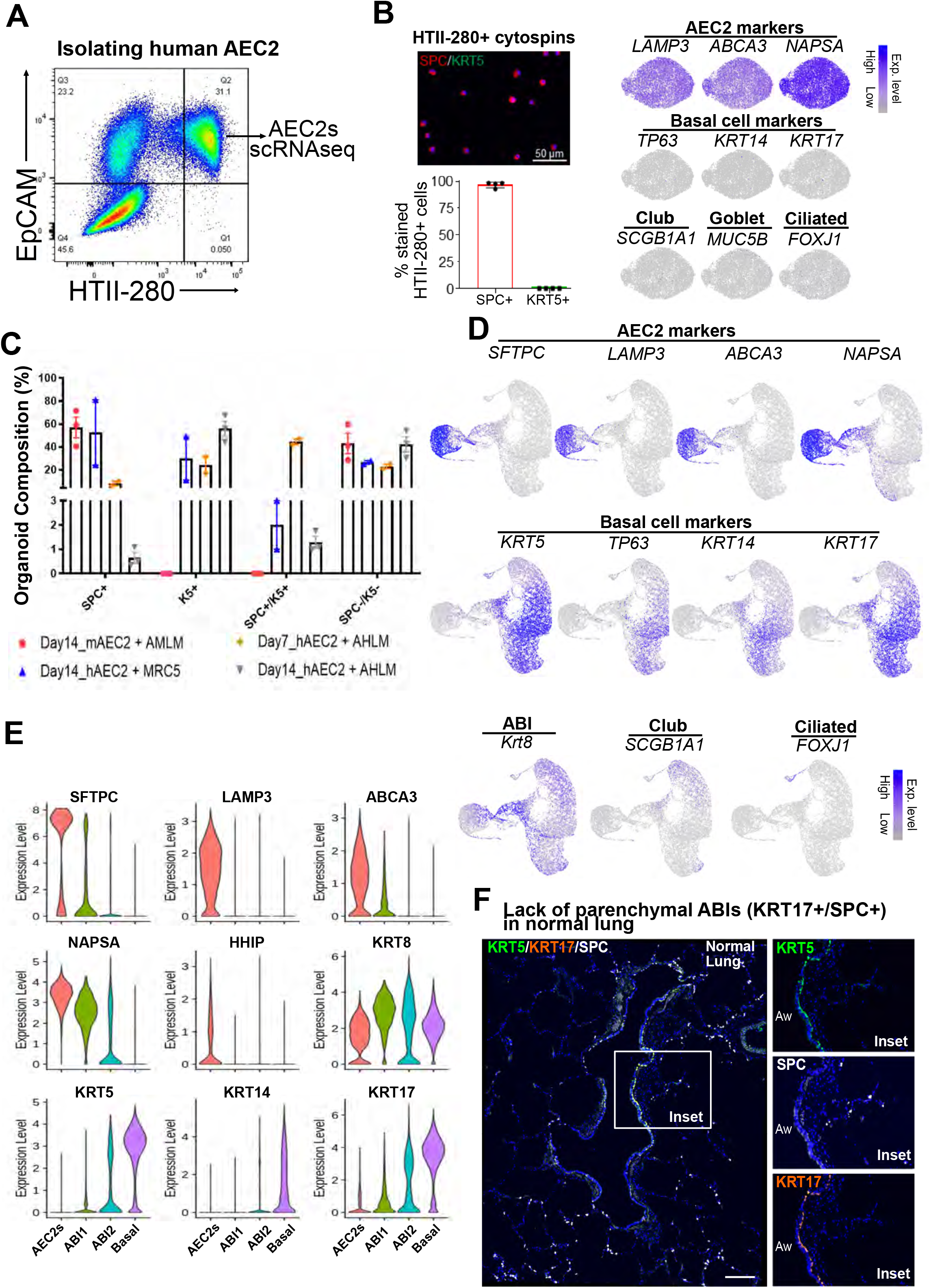
Isolation and differentiation of mature hAEC2s. (A) FACS plot shows the strategy for isolating hAEC2s. (B) Quantification of SFTPC+ and KRT5+ cells in freshly sorted HTII-280+ cells by cytospins and confirmation of the purity of HTII-280+ cells by scRNAseq. (C) Percentage of organoid composition in hAEC2-derived organoids co-cultured with AHLM at D7 and D14, in hAEC2-derived organoids co-cultured with MRC5 at D14, and in mAEC2-derived organoids co-cultured with AMLM at D14. (D) Gene feature plots show expression of hAEC2, basal cell, club, ciliated cell, and ABI markers in the merged UMAP plot of D0, D7, D14, and D21 organoids. (E) Violin plots show expression of hAEC2, basal cell, and ABI markers in different cell types. (F) Staining showing no evidence of KRT17+/SFTPC+ cells in the normal human lung.

**Figure S2 related to Figure 1.**
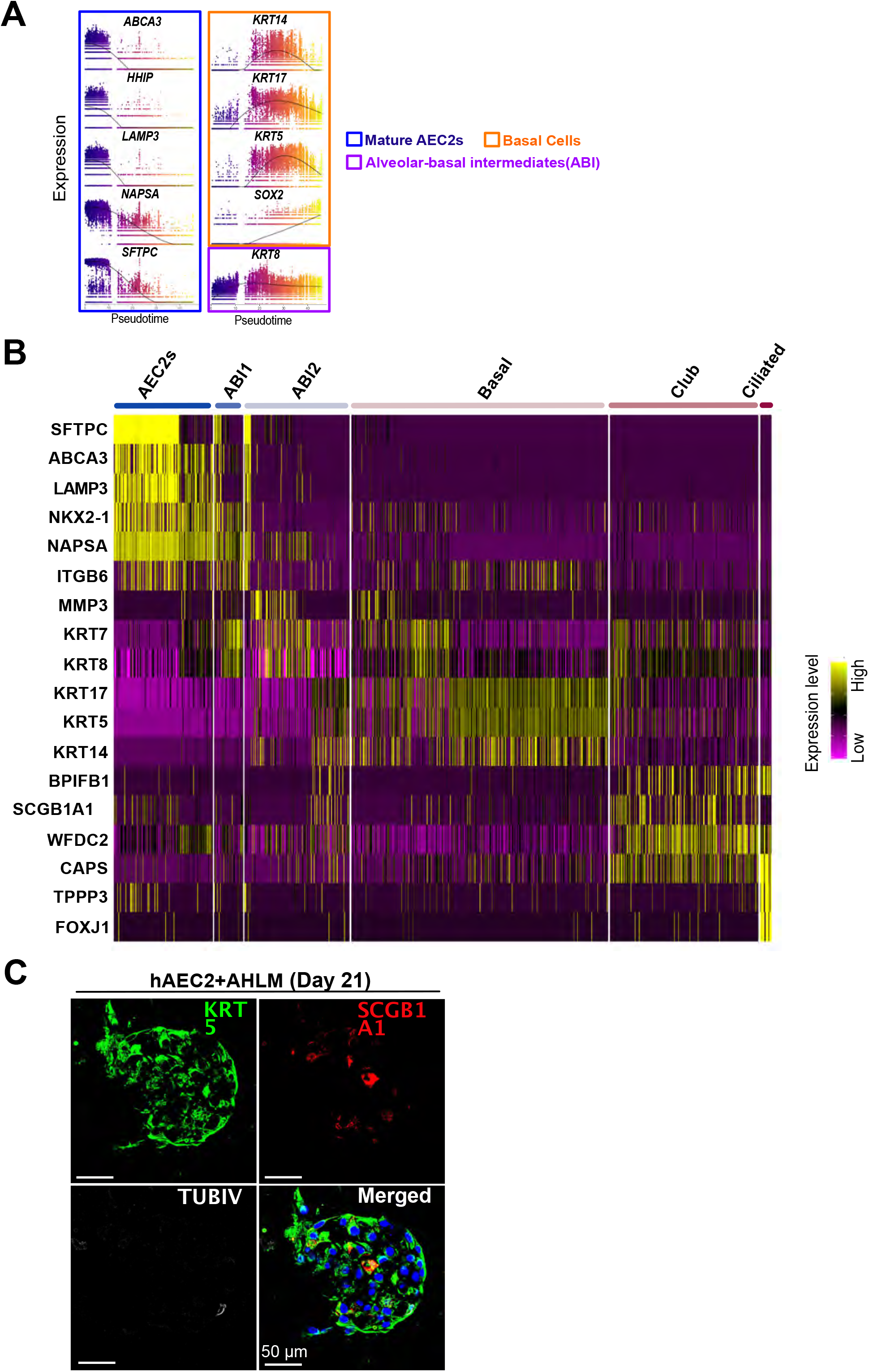
Expression profile of cellular subsets arising from hAEC2-derived organoids. (A) Expression along pseudotime trajectory of AEC2, ABI, and basal cell markers. (B) Heatmap of lineage marker expressions in different cellular subsets arising from hAEC2-derived organoids. (C) IHC of D21 organoids showing presence of SCGB1A1+ club cells and TUBIV+ ciliated cells.

**Figure S3 related to Figure 2.**
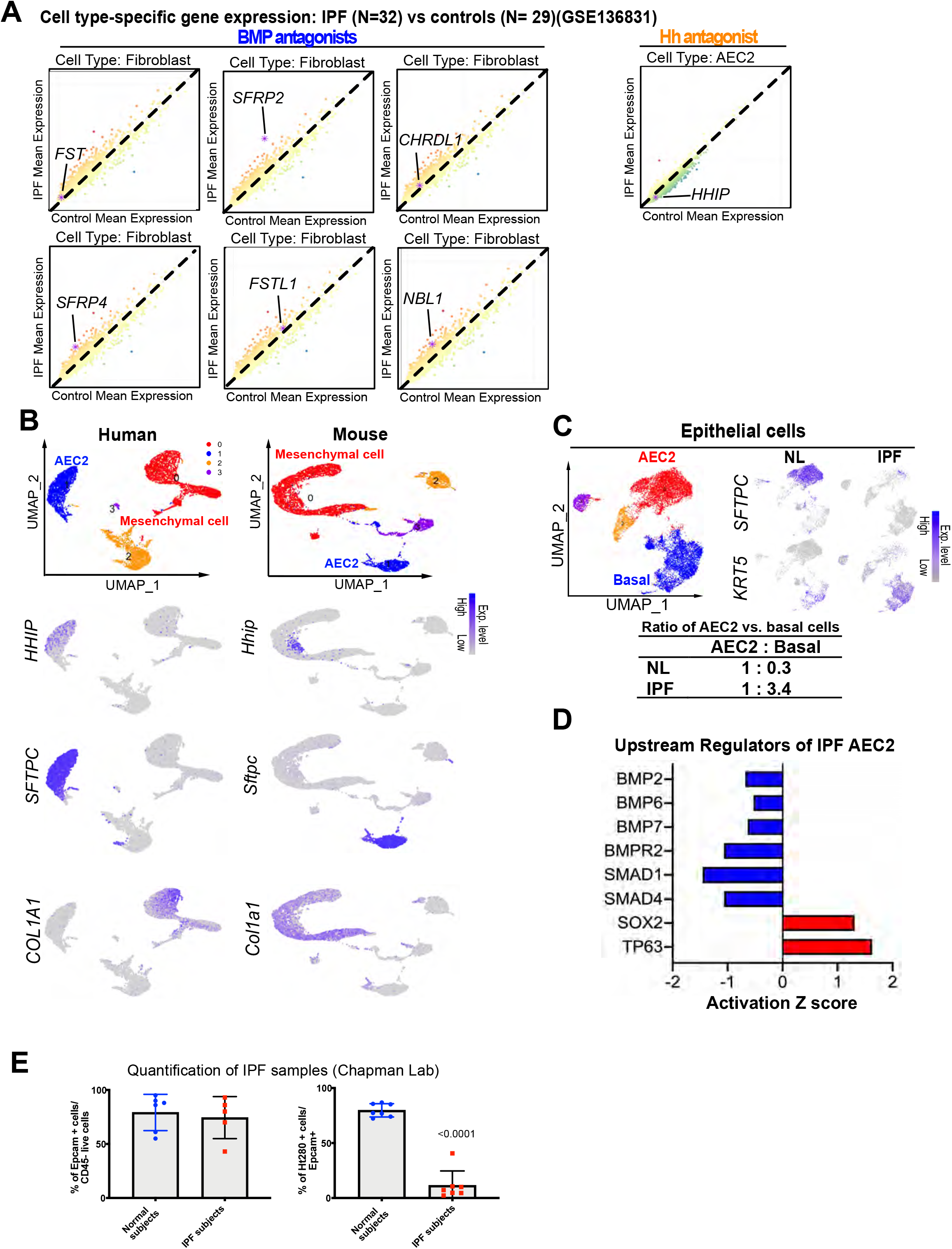
Dysregulation of hAEC2-intrinsic and niche factors in IPF. (A) Upregulation of BMP antagonists in IPF fibroblasts and downregulation of Hh antagonist, *HHIP*, in IPF AEC2s. (Data extracted from http://www.ipfcellatlas.com, GSE136831). (B) UMAP plots show that *HHIP* is predominantly expressed in SFTPC+ AEC2s in the human lung. In the mouse lung, *Hhip* is mainly expressed in COL1A1+ mesenchymal cells. (C) UMAP plots show less *SFTPC+* AEC2s but more *KRT5+* basal cells in IPF lungs (n = 2), compared to normal lungs (NL, n = 2). (D) IPA upstream regulator analysis shows downregulation of BMP activation and upregulation of basal cell differentiation pathways in IPF AEC2s, compared to NL AEC2s.. (E) Percentage of EpCAM+ cells/live cells and HTII-280+ cells/EpCAM+ cells of NL (n = 6) vs. IPF lungs (n = 7), quantified by FACS.

**Figure S4 related to Figure 3.**
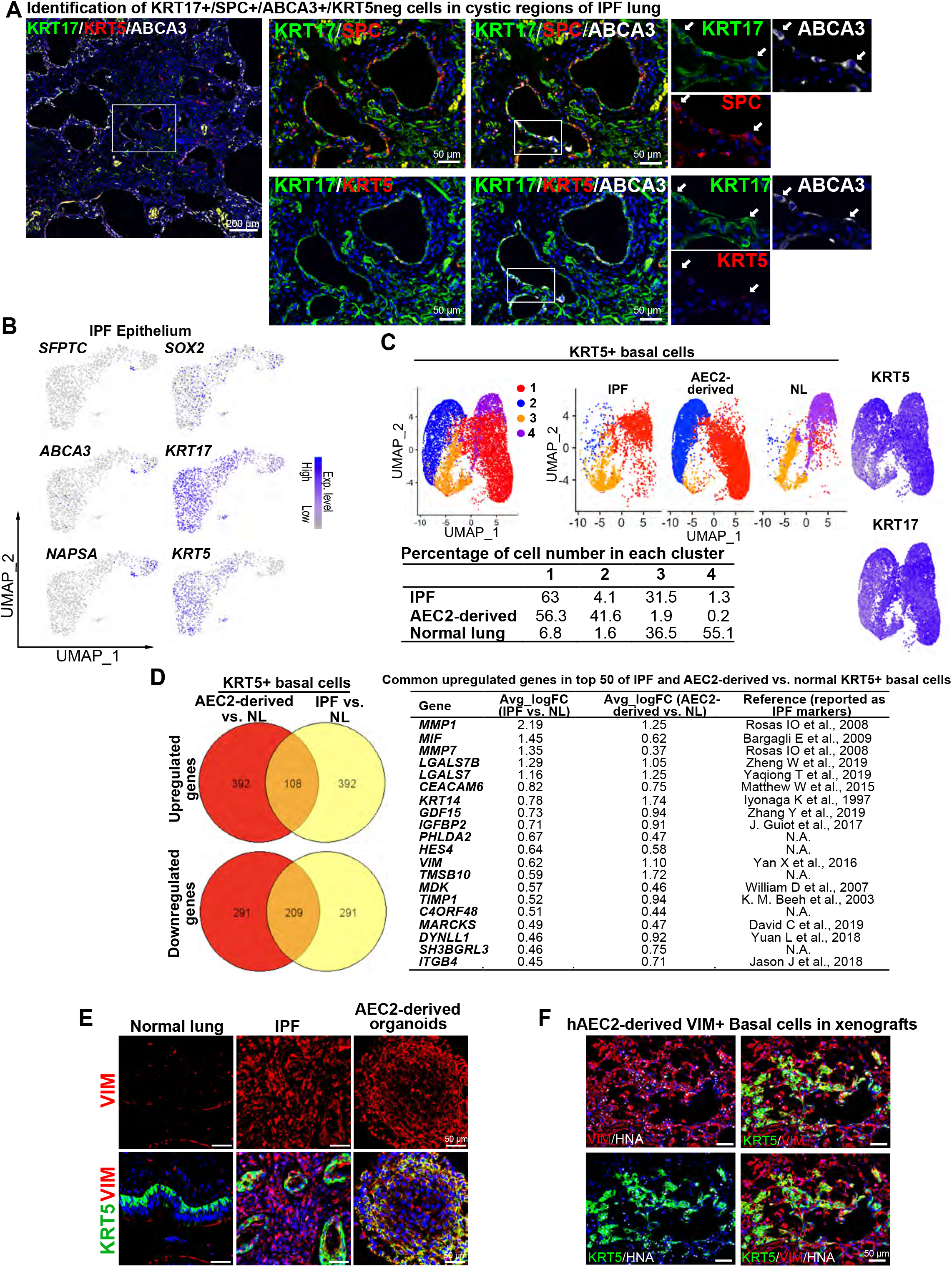
Identification of ABIs in the lungs of IPF patients and increased similarities between IPF and hAEC2-derived organoid basal cells. (A) Identification of KRT17+/SFTPC+/ABCA3+/KRT5-cells in cystic regions of IPF lung by immunostaining. (B) Gene feature plots show the expression of AEC2 markers and basal cell markers. (C) UMAP plots show *KRT5+* basal cells of IPF lungs (n = 2), hAEC2-deived organoids (D14 and D21), and NLs (n = 2). (D) GeneVenn plots show the overlapping of upregulated genes and downregulated genes in AEC2-derived basal cells and IPF basal cells against NL basal cells. Common upregulated genes in top 50 of IPF and AEC2-derived vs. normal KRT5+ basal cells are listed and most of them have been reported as IPF-associated markers. (E) Immunostaining of VIMENTIN (VIM) and KRT5 in NL, IPF lung, and hAEC2-derived organoids at D14. (F) Immunostaining of VIMENTIN (VIM), KRT5 and HNA in hAEC2-derived xenografts.

## STAR ★ METHODS

Detailed methods are provided in the online version of this paper and include the following:

- KEY RESOURCES TABLE
- CONTACT FOR REAGENT AND RESOURCE SHARING
- EXPERIMENTAL MODEL AND SUBJECT DETAILS

∘ Human Lung Tissue
∘ Animal Studies and Treatment
- METHOD DETAILS

∘ Lung Digestion and Fluorescence-activated Cell Sorting (FACS)
∘ Freezing and Thawing Primary Human Cells
∘ Cell Culture and Organoid Assay
∘ Quantitative RT-PCR (qPCR)
∘ Histology and Immunofluorescence
∘ Xenotransplantation
- QUANTIFICATION AND STATISTICAL ANALYSIS

∘ Statistical Analysis
∘ Immunofluorescence Image Quantification
∘ Single Cell RNA Sequencing Analysis
∘ Bulk RNA Sequencing Analysis
- DATA AND SOFTWARE AVAILABILITY

## STAR ★ METHODS

### KEY RESOURCES TABLE

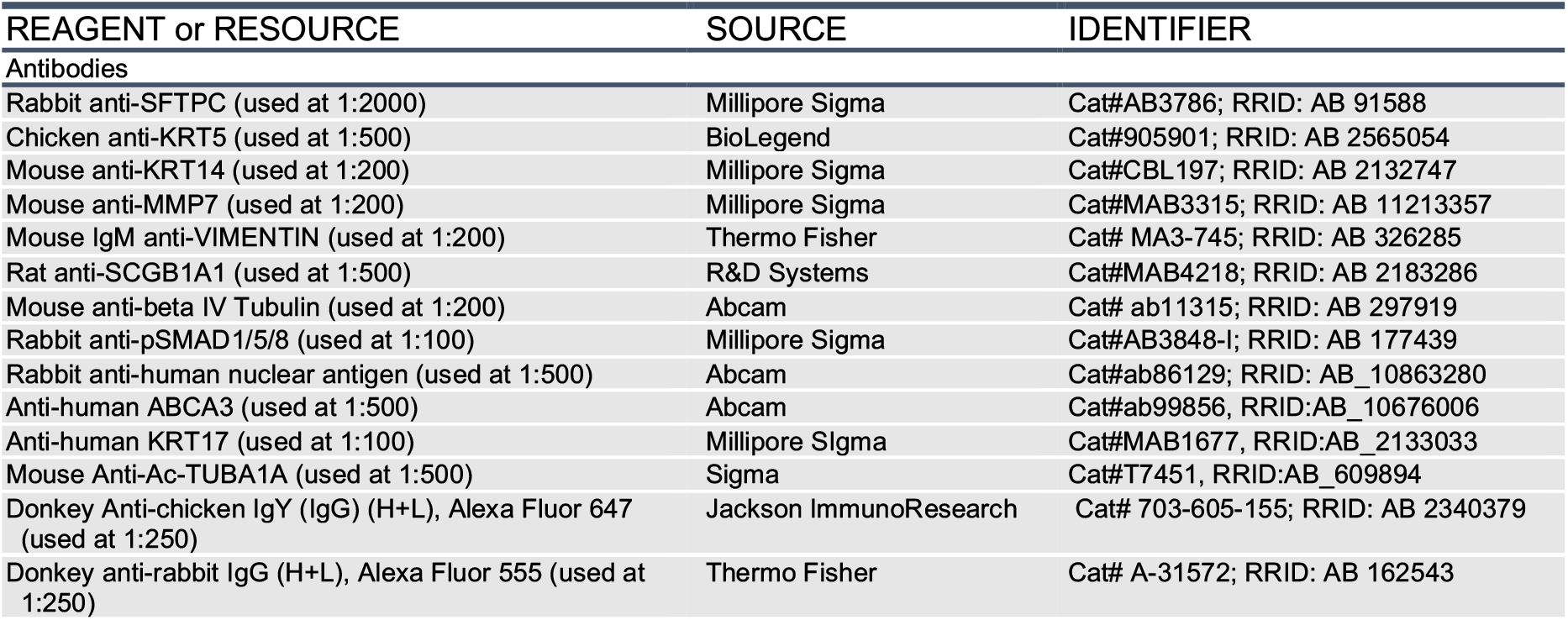
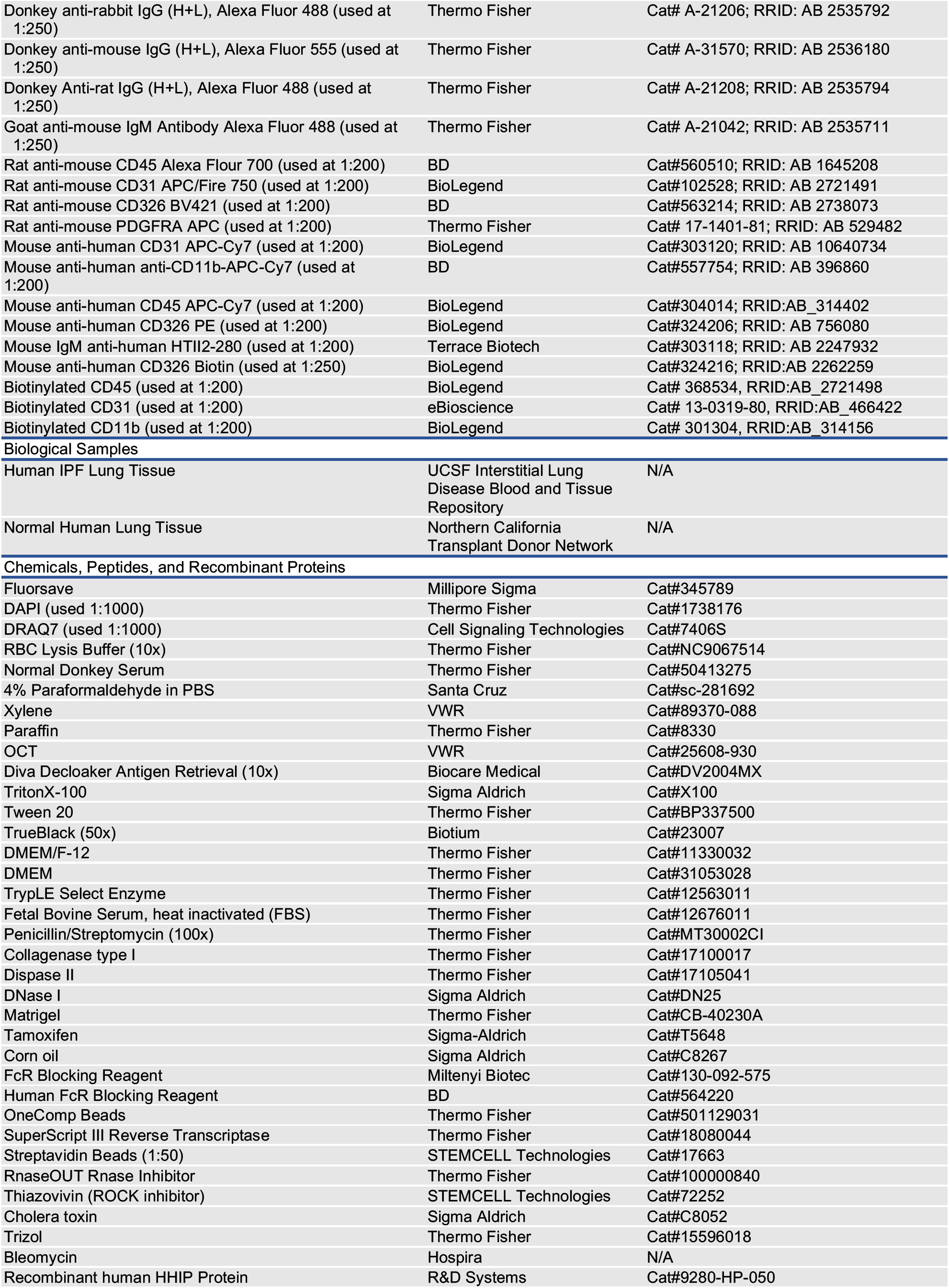
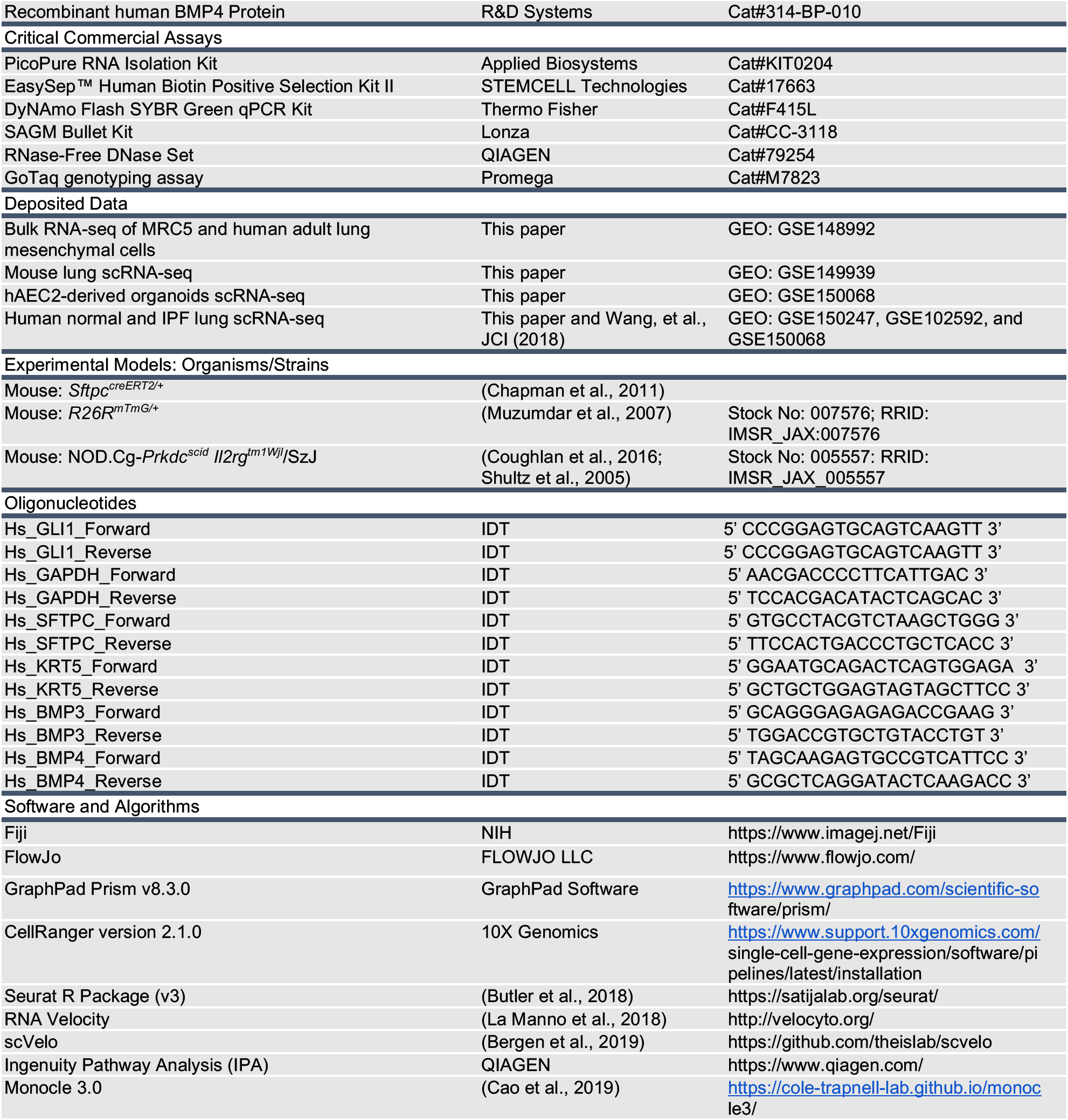

### CONTACT FOR REAGENT AND RESOURCE SHARING

Further information and requests for reagents may be directed to, and will be fulfilled by Hal Chapman and Tien Peng.

## EXPERIMENTAL MODEL AND SUBJECT DETAILS

### Human Lung Tissue

Studies involving human tissue were approved by the UCSF Institutional Review Board. IPF lungs were obtained from patients diagnosed with interstitial pneumonia or scleroderma at the time of lung transplant. In addition, tissues from diagnostic biopsies of ILD patients that proved to have UIP pattern were analyzed. All subjects provided written informed consent. Normal lungs were obtained from brain-dead donors that were rejected for lung transplantation.

### Animal Studies and Treatment

All animals were housed and treated in accordance with the Institutional Animal Care and Use Committee (IACUC) protocol approved at the University of California, San Francisco. Generation and genotyping of the *Sftpc^creERT2^ (Chapman et al., 2011)* and *R26R^mTmG^ (Muzumdar et al., 2007)* lines were performed as previously described. Animals between the ages of 8 and 12 weeks old were used for the experiments. For labelling mouse AEC2s (mAEC2s), tamoxifen (Cat#T5648; Sigma-Aldrich) was dissolved in corn oil and administered intraperitoneally at 200 mg/kg body weight per day for three consecutive days, followed by a week before mAEC2s were sorted. NOD.Cg-*Prkdc^scid^ Il2rg^tm1Wjl^/SzJ* (NOD *scid* gamma; NSG) mice were previously described (Coughlan et al., 2016; Shultz et al., 2005). Animal studies utilized a minimum of 4 mice per group. Mice were injured with oral aspiration of bleomycin (2.1 U/kg body weight). Mice were weighed twice a week. Mice were sacrificed between Day 17-20 post injury for histopathological analysis.

## METHOD DETAILS

### Histology and Immunofluorescence

#### Paraffin Embedding

The right ventricle of the mice were perfused with 1-3 mL PBS and the lungs were inflated with 1-3 mL 4% paraformaldehyde (PFA) in PBS, and then fixed in 4% PFA overnight at 4 °C. After fixation, the lungs were washed with cold PBS 4 times for 30 minutes each at 4 °C, and then dehydrated in a series of increasing ethanol concentration washes (30%, 50%, 70%, 95% and 100%) for a minimum of 2 hours per wash. The dehydrated lungs were incubated with xylene for 1 hr at RT and then with paraffin at 65 °C for 90 minutes twice, and then embedded in paraffin. The lungs were sectioned at 8 μm on a microtome.

#### OCT Embedding

Lung tissues of euthanized mice were processed as described before (Kathiriya et al., 2020). Briefly, 94%OCT/2%PFA/4%PBS-inflated lungs were fixed with 4% PFA for 1 hour at room temperature, washed with PBS for 4 hrs at room temperature, and embedded in OCT after 30% and 15% sucrose gradient washing. Organoids in 3D matrigel were fixed with 4% PFA for 30 minutes at room temperature or overnight at 4 °C, then washed in PBS overnight at least three times, followed by embedding in OCT. 8 μm sections were cut on a cryostat for both mouse lungs and organoids.

#### Immunofluorescent staining

Paraffin sections were incubated in xylene for 10 minutes twice, then rehydrated in ethanol washes (100% twice, 95%, 70%, 50% ethanol) for 5 minutes each. OCT embedded slides were fixed in 4% PFA at RT for 10 minutes, then washed three times with PBS. For both paraffin and OCT embedded slides, antigen retrieval (Cat#DV2004MX; Biocare Medical) was performed for 30 minutes at 95 °C or at 155 °C followed by incubation with sodium borohydride (Sigma) in PBS to reduce aldehyde-induced background fluorescence. Slides were washed with 0.1% Tween 20 in PBS (PBST) 3 times for 5 minutes. Slides were then incubated in blocking buffer (3% donkey serum in PBST) for at least 1 hour, then incubated overnight in primary antibodies in 1% donkey serum in PBST at 4 °C. The following primary antibodies were used for staining: rabbit anti-SFTPC (Cat#AB3786; Millipore Sigma; used 1:2000), chicken anti-KRT5 (Cat#905901; BioLegend; used at 1:500), Mouse anti-KRT14 (Cat#CBL197; Millipore Sigma; used at 1:200), mouse anti-MMP7 (Cat#MAB3315; Millipore Sigma; used at 1:200), mouse IgM anti-VIMENTIN (Thermo Fisher; Cat# MA3-745; used at 1:200), mouse anti-beta IV Tubulin (Abcam; Cat# ab11315; used at 1:200), mouse anti-Ac-Tuba1a (Sigma; Cat# T7451, used at 1:500), RRID rabbit anti-p-SMAD1/5/8 (Cat#AB3848-I; Millipore Sigma; used 1:100), and rat anti-SCGB1A1 (R&D; Cat#MAB4218; used at 1:500). Slides were washed with PBST and then incubated for 1 hour at RT in secondary antibodies diluted in PBST. The following secondary antibodies were used at 1:250: Donkey Anti-chicken IgY (IgG) (H+L) Alexa Fluor 647 (Jackson ImmunoResearch; Cat# 703-605-155; used at 1:250), Donkey anti-rabbit IgG (H+L) Alexa Fluor 555 (Thermo Fisher; Cat# A-31572; used at 1:250), Donkey anti-rabbit IgG (H+L) Alexa Fluor 488 (Thermo Fisher; Cat# A-21206; used at 1:250), Donkey antimouse IgG (H+L) Alexa Fluor 555 (Thermo Fisher; Cat# A-31570; used at 1:250), Donkey Anti-rat IgG (H+L) Alexa Fluor 488 (Thermo Fisher; Cat# A-21208; used at 1:250), and Goat anti-mouse IgM Antibody Alexa Fluor 488 (Thermo Fisher; Cat# A-21042; used at 1:250). DAPI (0.2 μg/mL) (Cat#1738176; Thermo Fisher) was added for 5 minutes, and slides were then mounted with TrueBlack to reduce background and Fluorosave to maintain fluorescence. Images were captured using Zeiss Imager M1 and analyzed using Axiovision 4.8.2 (Zeiss, Germany). Where indicated, multiple images at 20X were captured using the “MosaiX” function and stitched together using “Tile Stitch” function in the Axiovision software.

### Lung digestion and Fluorescence Activated Cell Sorting (FACS)

Human lung tissue was obtained and dissected into 5 cm^3^ pieces. Tissue was washed two times in 500 ml sterile PBS and one time in HBSS for 10 min at 4 °C. Using autoclaved Kim Wipes, tissue was compressed to remove as much liquid as possible and further dissected into 1 cm^3^ pieces. Sterile HBSS buffer containing 15 U/ml Dispase II (Cat#17105041; Thermo fisher), 225 U/mL Collagenase Type I (Cat#17100017; Thermo Fisher), 100 U/mL Dnase I (Cat#DN25; Sigma-Aldrich), and 1% penicillin/streptomycin was added to the tissue pieces. Tissue was digested for 2 h at 37 °C and Fungizone (1:400) was added for the final 30 min of the digest. After digest the enzyme solution was diluted with an equal volume of sort buffer (DMEM without phenol red, 2% FBS and penicillin/streptomycin) and liquefied using an Osterizer 12 Speed Blender as follows: 5 s pulse, 5 s grate, and 5 s pulse. The suspension was poured through a glass funnel lined with sterile 4×4 gauze, applying some compression to recover as much of the solution as possible. The cell suspension was sequentially filtered through 100um, 70um and 40umstrainers. Red blood cells were removed using Red Blood Cell Lysis Buffer (Sigma-Aldrich). Immune cells were depleted by resuspending 2.0 × 10^7^ cells/ml and FC blocked for 10 minutes rocking at 4 °C. Biotinylated CD45 (clone 2D1), CD31 (clone WM-59), and CD11b (clone ICRF44) were spiked into the cell suspension at 1:200 and rocked an additional 20 minutes. Streptavidin beads (Cat#17663; STEMCELL Technologies; used 1:50) were added at 25ul/ml to exclude immune and endothelial cells using MagRack 6 (GE Healthcare 89129-096). Cells were either frozen for future use or stained for flow cytometry. The following FACS antibodies were used at 1:200: mouse anti-human CD45 APC-Cy7 (BioLegend; Cat#304014; used at 1:200), mouse anti-human anti-CD11b-APC-Cy7 (BD; Cat#557754; used at 1:200), mouse anti-human CD31 APC-Cy7 (BioLegend; Cat#303120; used at 1:200), mouse anti-human CD326 PE (BioLegend; Cat#324206; used at 1:200), and mouse IgM anti-human HTII-280 (Terrace Biotech, Cat#303118; used at 1:200). After 30 min incubation, goat anti-mouse IgM Alexa Fluor 488 (Thermo Fisher; Cat# A-21042; used at 1:1000) was added to label HTII-280+ cells. Immune and endothelial cells were further excluded on FACS by gating CD45-/CD11b-/CD31-. Human AEC2s were sorted for EpCAM+/HTII-280+ cells, and adult human lung mesenchymal cells (AHLM) were sorted for CD45-/CD11b-/CD31-/EpCAM-cells.

For mouse, whole lung was dissected from adult animals and perfused with a digestion cocktail of Collagenase Type I (Cat#17100017; Thermo Fisher; used 225 U/mL), Dispase II (Cat#17105041; Thermo fisher; used 15 U/mL) and Dnase I (Cat#DN25; Sigma-Aldrich; used 50 U/mL) and removed from the chest. The lung was further diced with a razor blade and incubated in the digestion cocktail for 45 mins at 37 °C with continuous shaking. The mixture was then washed with sorting buffer (2% FBS and 1% Penicillin-Streptomycin in DMEM (Cat#31053028; Thermo Fisher)). The mixture was passed through a 70 μm cell strainer and resuspended in RBC lysis buffer (Cat#NC9067514; Thermo Fisher), then passed through a 40 μm cell strainer. Cell suspensions were incubated with the appropriate conjugated antibodies in sorting buffer for 30 min at 4 °C and washed with sorting buffer. Doublets and dead cells were excluded based on forward and side scatters and DRAQ7 (Cat#7406S; Cell Signaling Technologies) fluorescence, respectively. Immune and endothelial cells were excluded using CD45 (Alexa Flour 700; Cat#560510; BD; used 1:200) and CD31 (APC/Fire750; Cat#102528; BioLegend; used 1:200), respectively. Mouse AEC2s (mAEC2s) were sorted using endogenous GFP from tamoxifen-induced adult *Sftpc^creERT2/+^:R26^mTmG/+^* lungs. Mouse lung mesenchymal cells (AMLM) were sorted from adult wild-type mouse lung by excluding immune, endothelial cells, and epithelial cells (anti-CD326 BV421; Cat#563214; BD; used at 1:200), based on the selection of EpCAM-/CD45-/CD31-. Cells were sorted into sorting buffer. Analysis was performed using FlowJo software.

### Freezing/Thawing primary human cells

To freeze, prior to isolation a 2x Freeze Solution was prepared, filtered, and stored at 4C (2% 1.5 M Hepes, 10% FBS, 78% F12 and 10% DMSO). Cells and all solutions were kept on ice during this protocol. Cells were resuspended in F12 media and counted. Volume was adjusted to a density of 10^7 cells/mL, followed by slow addition of equal volume of 2x Freeze Solution to the cell suspension with gentle mixing. Desired volume of cell aliquots was then frozen and stored in liquid nitrogen. To thaw, DMEM with 10% FBS was warmed to 37C. Cryovials were thawed in 37C water bath and cells were transferred to a 15mL centrifuge tube. The cell suspension was slowly diluted by adding an equal volume of warm media and incubated for one minute, followed by addition of 12 mL of warmed DMEM to the centrifuge tube. Cells were spun at 550g for 4 minutes and stained for flow cytometry.

### Cell Culture and Organoid Assay

MRC5 cells were purchased from ATCC (cat#CCL-171). All the mesenchymal cells (AHLM, AMLM, and MRC5) were cultured in DMEM/F-12 (Cat#11330032; Thermo Fisher) with 10% FBS and 1% Penicillin-Streptomycin. Media was refreshed every other day and primary lung mesenchymal cells were maintained for no more than five passages.

For organoids assay, AEC2 and mesenchymal cells were co-cultured (5,000 AEC2:30,000 mesenchymal cells/well) in a modified MTEC media diluted 1:1 in growth factor-reduced matrigel (Cat#CB-40230A; Thermo Fisher). Modified MTEC culture media is comprised of small airway basal media (SABM, Cat#CC-3118; Lonza) with selected components from SAGM bullet kit including Insulin, Transferrin, Bovine Pituitary Extract, Retinoic Acid, and human Epidermal Growth Factor. 0.1 μg/mL cholera toxin (Cat#C8052; Sigma Aldrich), 5% FBS, and 1% Penicillin-Streptomycin were also added to the media. Cell suspension-matrigel mixture was placed in a transwell and incubated in growth media with 10 μM ROCK inhibitor (Cat#72252; STEMCELL Technologies) in a 24 well plate for 48 hours, after which the media was replenished every other day (lacking ROCK inhibitor). Each experimental condition was performed in triplicates. Where applicable, recombinant BMP4 (Cat#314-BP-010; R&D Systems; used 50 ng/mL) and HHIP (Cat#9280-HP-050; R&D Systems; used 5 μg/mL) were added to the media after 48 hours and replenished in every media change. Colonies were assayed after 7, 14, and 21 days.

To extract RNA from the organoid assays, cell suspension-matrigel mixtures in the transwells were washed with PBS and incubated in a digestion cocktail of Collagenase Type I (Cat#17100017; Thermo Fisher; used 225 U/mL), Dispase II (Cat#17105041; Thermo fisher; used 15 U/mL), and Dnase I (Cat#DN25; Sigma-Aldrich; used 50 U/mL) for 1 hour at 37 °C with intermittent resuspension. The mixture was removed from the transwell and resuspended in TrypLE (Cat#12563011; Thermo Fisher) and shaken at 37 °C for 20 minutes. Cells were resuspended in sorting buffer (2% FBS and 1% Penicillin-Streptomycin in DMEM) and blocked with human FcR blocking reagent (Cat#564220; BD; used 1:50) for 10 minutes at 4 °C. Cells were then stained with biotin anti-human CD326 (Cat#324216; BioLegend; 1:250) for 30 minutes at 4 °C. Streptavidin beads (Cat#17663; STEMCELL Technologies; used 1:50) were added to isolate the epithelial cells using MagRack 6 (GE Healthcare 89129-096), and the rest cells in the suspension were mesenchymal cells. For sequencing of cells from organoids, organoids were digested as above and FACS sorted for live EpCAM+ cells.

To analyze the organoids by histology, organoids in the transwell were washed with PBS and fixed with 4% PFA overnight at 4 °C. After fixation, 30% sucrose was added in the organoids. After overnight incubation of 30% sucrose, the organoids were embedded in OCT and then sectioned.

### Quantitative RT-PCR

Total RNA was obtained from cells isolated from organoid assays using the PicoPure RNA Isolation Kit (Cat#KIT0204; Applied Biosystems), following the manufacturers’ protocols. cDNA was synthesized from total RNA using the SuperScript Strand Synthesis System (Cat#18080044, Cat#100000840; Thermo Fisher). Quantitative RT-PCR (qRT-PCR) was performed using the SYBR Green system (Cat#F415L; Thermo Fisher). Primers are listed in KEY RESOURCES TABLE. Relative gene expression levels after qRT-PCR were defined using the ΔΔCt method and normalizing to GAPDH

### Xenotransplantation Assay

500,000 freshly sorted hAEC2s (as live/CD45neg/EpCAM+/HTII-280+) in 40uL volume (1XPBS) from normal donor lungs were transplanted at 10 days post bleomycin injury into the lungs of NOD *scid* gamma mice via oral aspiration as described previously (Kathiriya et al., 2020). For co-transplantation, 500,000 hAEC2s were mixed with 200,000 primary AHLMs in 40 uL volume (1X PBS) and delivered into the lungs of injured mice via oral aspiration. Three mice in each group were analyzed for with at least two sections taken at 200μm apart to capture different regions of the lung. Any HNA+ (human nuclear antigen) region of >5 cells was counted as one engrafted region. At least three lobes were analyzed at each section for three mice/condition. Data are presented as mean ± SD. ** p < 0.01 (Student’s t-test).

### Single cell transcriptomics

Single cell sequencing was performed on a 10X Chromium instrument (10X Genomics, Pleasanton, CA) at the Institute of Human Genetics (UCSF, San Francisco, CA) as described before. Briefly, cells were isolated via flow cytometry according to the experimental design, washed twice, and re-suspended in 1X PBS supplemented with 0.04% BSA at 1000 cells/μL. Single cells were then loaded on a Chromium Controller instrument (10X Genomics) to generate single-cell Gel Bead-In-EMulsions (GEMs), which were used for library preparation. Libraries were prepared by performing reverse transcription on Bio-Rad C1000 Touch Thermal Cycler (Bio-Rad, Hercules, CA), following which, GEMs were harvested to amplify cDNA also using Bio-Rad C1000 Touch Thermal Cycler. SPRIselect (Beckman Coulter, Brea, CA) was used to select for amplified cDNA. Indexed sequencing libraries were constructed using the Chromium Single-Cell 3’ library kit (10X Genomics) and sequenced on NovaSeq 6000 (Illumina) with the following parameters: Read 1 (26 cycles), Read 2 (98 cycles), and i7 index (8 cycles) to obtain a sequencing depth of ~100,000 read/cell. Reads were aligned to appropriate mouse or human genome and quantified using Cell Range Single-Cell Software Suite.

## QUANTIFICATION AND STATISTICAL ANALYSIS

### Statistical Analysis

All statistical analyses were performed in GraphPad Prism. Unpaired one-tailed t-tests were used to determine the p-value and data in graphs are presented as mean ± SEM or mean ± SD.

### Immunofluorescence Image Quantification

Sections were imaged for quantification on a Zeiss Lumar V12 microscope. At least three samples per genotype/condition were used, and at least 4 randomly selected sections were chosen for each sample. Cell counts for stained cells were performed on Fiji using the “Cell Counter” plug-in. Cell counts for DAPI were performed automatically by Fiji software using the “Analyze Particles” tool. At least three samples per condition were used, and at least 6 randomly selected sections were chosen for each sample. For all analyses, the performers were blinded to the specimen genotype and condition. Results were averaged between each specimen and SEM or SD were calculated per condition.

### Analysis of single cell mRNA-sequencing

To build transcript profiles of individual cells the CellRanger version 2.1.1 software with default settings was used for de-multiplexing, aligning reads with STAR software to GRCm38 for mouse genome and hg19 or GRCh38 for human, and counting unique molecular identifiers (UMIs). Pre-processing data, integration, clustering, and Uniform Manifold Approximation and Projection (UMAP) was performed in Seurat v3 package (Stuart et al., 2019). In total, we filtered the data in 2 different steps. We first filtered the dataset by only accepting cells that expressed a minimum of 200 genes and genes that were expressed in at least 3 cells. Our second filter was set to accept cells with less than 6000 unique gene counts and 10% mitochondrial counts. After removing unwanted cells, we used “LogNormalize” to normalize the gene expression measurements for each cell. We calculated 2,000 features that exhibit high cell-to-cell variation, which were used in principle component analysis (PCA) after scaling the data. We used the JackStrawPlot function in the Seurat package to create Scree plots and compare p-value (significance) for each principle component. We selected 10 different PCA’s for clustering of both mouse and human cells. Clustering results were visualized using the UMAP algorithm in the Seurat package. For cluster visualization and individual gene visualization on all clusters we used the UMAP function. IPA analysis was performed using the differentially expressed genes with adj. p <0.1 and logfc >0.15 or <-0.15.

### Trajectory and Velocity analysis

Trajectory analysis was performed using Moncole 3 (Cao et al., 2019) by importing the counts from the Seurat integrated object and RNA velocity was performed using scVelo (Bergen et al., 2019). Briefly, the aligned bam files were used as input for Velocyto (La Manno et al., 2018) to calculate unspliced and spliced reads and generate loom files, which were read into Seurat using ReadVelocity() function in SeuratWrappers and converted into Seurat object using as.Seurat() function. Following processing individual samples to filter out low quality/dying cells as described before, samples were integrated and visualized via UMAP. After manual curation of cell types, six major cell types were identified. Counts of the Seurat integrated object were imported into Monocle 3 to infer trajectory. AEC2s from normal lung (day 0) were called as root node to infer pseudotime trajectory. We employed scVelo to predict future states of individual cells. The normalized and log transformed data with UMAP coordinates and cluster identities from Seurat was used to estimate velocities and velocity graph was constructed using scvelo.tl.velocity(), with stochastic mode, and scvelo.tl.velocity_graph().

### Bulk RNA Sequencing Analysis

Total RNA was extracted from AHLM and MRC5 cells using the the RNeasy Kit (Cat#74004; QIAGEN), following the manufacturers’ protocols. Sequencing was done using HiSeq 4000. Quality control of reads was conducted by using FastQC (Babraham Bioinformatics). Ligation adaptors were removed using the Cutadapt and Sickle. Sequencing reads were aligned using STAR and UCSC human GRCh38/hg38 was used as reference genome. The differential gene expression list was generated using DEseq.

## DATA AND SOFTWARE AVAILABILITY

The mouse scRNA-seq data and human bulk RNA-seq data reported in this paper is deposited in NCBI Gene Expression Omnibus (GEO) under the accession number: GSE149939 and GSE148992, respectively. The human scRNA-seq data from normal, IPF, and organoids reported in this paper is deposited in GEO under the following accession numbers: GSE150068 and GSE150247.

## Notes

### Competing Interest Statement

The authors have declared no competing interest.

## REFERENCES

Adams, T.S., Schupp, J.C., Poli, S., Ayaub, E.A., Neumark, N., Ahangari, F., Chu, S.G., Raby, B., DeIuliis, G., Januszyk, M., et al. (2019). Single Cell RNA-seq reveals ectopic and aberrant lung resident cell populations in Idiopathic Pulmonary Fibrosis. bioRxiv.

Barkauskas, C.E., Cronce, M.J., Rackley, C.R., Bowie, E.J., Keene, D.R., Stripp, B.R., Randell, S.H., Noble, P.W., and Hogan, B.L. (2013). Type 2 alveolar cells are stem cells in adult lung. J Clin Invest 123, 3025–3036.

Bergen, V., Lange, M., Peidli, S., Wolf, F.A., and Theis, F.J. (2019). Generalizing RNA velocity to transient cell states through dynamical modeling. bioRxiv.

Butler, A., Hoffman, P., Smibert, P., Papalexi, E., and Satija, R. (2018). Integrating single-cell transcriptomic data across different conditions, technologies, and species. Nat Biotechnol 36, 411–420.

Cao, J., Spielmann, M., Qiu, X., Huang, X., Ibrahim, D.M., Hill, A.J., Zhang, F., Mundlos, S., Christiansen, L., Steemers, F.J., et al. (2019). The single-cell transcriptional landscape of mammalian organogenesis. Nature 566, 496–502.

Cassandras, M., Wang, C., Kathiriya, J., Tsukui, T., Matatia, P., Matthay, M., Wolters, P., Molofsky, A., Sheppard, D., Chapman, H., et al. (2019). <em>Gli1</em>+ mesenchymal stromal cells modulate epithelial metaplasia in lung fibrosis. bioRxiv.

Chapman, H.A., Li, X., Alexander, J.P., Brumwell, A., Lorizio, W., Tan, K., Sonnenberg, A., Wei, Y., and Vu, T.H. (2011). Integrin alpha6beta4 identifies an adult distal lung epithelial population with regenerative potential in mice. J Clin Invest 121, 2855–2862.

Chuang, P.T., Kawcak, T., and McMahon, A.P. (2003). Feedback control of mammalian Hedgehog signaling by the Hedgehog-binding protein, Hip1, modulates Fgf signaling during branching morphogenesis of the lung. Genes Dev 17, 342–347.

Coughlan, A.M., Harmon, C., Whelan, S., O’Brien, E.C., O’Reilly, V.P., Crotty, P., Kelly, P., Ryan, M., Hickey, F.B., O’Farrelly, C, et al. (2016). Myeloid Engraftment in Humanized Mice: Impact of Granulocyte-Colony Stimulating Factor Treatment and Transgenic Mouse Strain. Stem Cells Dev 25, 530–541.

Habermann, A.C., Gutierrez, A.J., Bui, L.T., Yahn, S.L., Winters, N.I., Calvi, C.L., Peter, L., Chung, M.-I., Taylor, C.J., Jetter, C., et al. (2019). Single-cell RNA-sequencing reveals profibrotic roles of distinct epithelial and mesenchymal lineages in pulmonary fibrosis. bioRxiv.

Jiang, P., Gil de Rubio, R., Hrycaj, S.M., Gurczynski, S.J., Riemondy, K.A., Moore, B.B., Omary, M.B., Ridge, K.M., and Zemans, R.L. (2020). Ineffectual AEC2-to-AEC1 Differentiation in IPF: Persistence of KRT8(hi) Transitional State. Am J Respir Crit Care Med.

Kathiriya, J.J., Brumwell, A.N., Jackson, J.R., Tang, X., and Chapman, H.A. (2020). Distinct Airway Epithelial Stem Cells Hide among Club Cells but Mobilize to Promote Alveolar Regeneration. Cell Stem Cell 26, 346–358 e344.

Kobayashi, Y., Tata, A., Konkimalla, A., Katsura, H., Lee, R.F., Ou, J., Banovich, N.E., Kropski, J.A., and Tata, P.R. (2019). Persistence of a novel regeneration-associated transitional cell state in pulmonary fibrosis. bioRxiv.

Koli, K., Myllarniemi, M., Vuorinen, K., Salmenkivi, K., Ryynanen, M.J., Kinnula, V.L., and Keski-Oja, J. (2006). Bone morphogenetic protein-4 inhibitor gremlin is overexpressed in idiopathic pulmonary fibrosis. Am J Pathol 169, 61–71.

Kumar, P.A., Hu, Y., Yamamoto, Y., Hoe, N.B., Wei, T.S., Mu, D., Sun, Y., Joo, L.S., Dagher, R., Zielonka, E.M., et al. (2011). Distal airway stem cells yield alveoli in vitro and during lung regeneration following H1N1 influenza infection. Cell 147, 525–538.

La Manno, G., Soldatov, R., Zeisel, A., Braun, E., Hochgerner, H., Petukhov, V., Lidschreiber, K., Kastriti, M.E., Lonnerberg, P., Furlan, A., et al. (2018). RNA velocity of single cells. Nature 560, 494–498.

Li, Y., Wu, Q., Sun, X., Shen, J., and Chen, H. (2020). Organoids as a Powerful Model for Respiratory Diseases. Stem Cells Int 2020, 5847876.

Muzumdar, M.D., Tasic, B., Miyamichi, K., Li, L., and Luo, L. (2007). A global double-fluorescent Cre reporter mouse. Genesis 45, 593–605.

Myllarniemi, M., Lindholm, P., Ryynanen, M.J., Kliment, C.R., Salmenkivi, K., Keski-Oja, J., Kinnula, V.L., Oury, T.D., and Koli, K. (2008). Gremlin-mediated decrease in bone morphogenetic protein signaling promotes pulmonary fibrosis. Am J Respir Crit Care Med 177, 321–329.

Prasse, A., Binder, H., Schupp, J.C., Kayser, G., Bargagli, E., Jaeger, B., Hess, M., Rittinghausen, S., Vuga, L., Lynn, H., et al. (2019). BAL Cell Gene Expression Is Indicative of Outcome and Airway Basal Cell Involvement in Idiopathic Pulmonary Fibrosis. Am J Respir Crit Care Med 199, 622–630.

Ray, S., Chiba, N., Yao, C., Guan, X., McConnell, A.M., Brockway, B., Que, L., McQualter, J.L., and Stripp, B.R. (2016). Rare SOX2(+) Airway Progenitor Cells Generate KRT5(+) Cells that Repopulate Damaged Alveolar Parenchyma following Influenza Virus Infection. Stem Cell Reports 7, 817–825.

Rock, J.R., Barkauskas, C.E., Cronce, M.J., Xue, Y., Harris, J.R., Liang, J., Noble, P.W., and Hogan, B.L. (2011). Multiple stromal populations contribute to pulmonary fibrosis without evidence for epithelial to mesenchymal transition. Proc Natl Acad Sci U S A 108, E1475–1483.

Rosas, I.O., Richards, T.J., Konishi, K., Zhang, Y., Gibson, K., Lokshin, A.E., Lindell, K.O., Cisneros, J., Macdonald, S.D., Pardo, A., et al. (2008). MMP1 and MMP7 as potential peripheral blood biomarkers in idiopathic pulmonary fibrosis. PLoS Med 5, e93.

Shultz, L.D., Lyons, B.L., Burzenski, L.M., Gott, B., Chen, X., Chaleff, S., Kotb, M., Gillies, S.D., King, M., Mangada, J., et al. (2005). Human lymphoid and myeloid cell development in NOD/LtSz-scid IL2R gamma null mice engrafted with mobilized human hemopoietic stem cells. J Immunol 174, 6477–6489.

Smirnova, N.F., Schamberger, A.C., Nayakanti, S., Hatz, R., Behr, J., and Eickelberg, O. (2016). Detection and quantification of epithelial progenitor cell populations in human healthy and IPF lungs. Respir Res 17, 83.

Strunz, M., Simon, L.M., Ansari, M., Mattner, L.F., Angelidis, I., Mayr, C.H., Kathiriya, J., Yee, M., Ogar, P., Sengupta, A., et al. (2019). Longitudinal single cell transcriptomics reveals Krt8+ alveolar epithelial progenitors in lung regeneration. bioRxiv.

Stuart, T., Butler, A., Hoffman, P., Hafemeister, C., Papalexi, E., Mauck, W.M., 3rd, Hao, Y., Stoeckius, M., Smibert, P., and Satija, R. (2019). Comprehensive Integration of Single-Cell Data. Cell 177, 1888–1902 e1821.

Wang, C., de Mochel, N.S.R., Christenson, S.A., Cassandras, M., Moon, R., Brumwell, A.N., Byrnes, L.E., Li, A., Yokosaki, Y., Shan, P., et al. (2018). Expansion of hedgehog disrupts mesenchymal identity and induces emphysema phenotype. J Clin Invest 128, 4343–4358.

Xi, Y., Kim, T., Brumwell, A.N., Driver, I.H., Wei, Y., Tan, V., Jackson, J.R., Xu, J., Lee, D.K., Gotts, J.E., et al. (2017). Local lung hypoxia determines epithelial fate decisions during alveolar regeneration. Nat Cell Biol 19, 904–914.

Xu, Y., Mizuno, T., Sridharan, A., Du, Y., Guo, M., Tang, J., Wikenheiser-Brokamp, K.A., Perl, A.T., Funari, V.A., Gokey, J.J., et al. (2016). Single-cell RNA sequencing identifies diverse roles of epithelial cells in idiopathic pulmonary fibrosis. JCI Insight 1, e90558.

Yang, Y., Riccio, P., Schotsaert, M., Mori, M., Lu, J., Lee, D.K., Garcia-Sastre, A., Xu, J., and Cardoso, W.V. (2018). Spatial-Temporal Lineage Restrictions of Embryonic p63(+) Progenitors Establish Distinct Stem Cell Pools in Adult Airways. Dev Cell 44, 752–761 e754.

Zacharias, W.J., Frank, D.B., Zepp, J.A., Morley, M.P., Alkhaleel, F.A., Kong, J., Zhou, S., Cantu, E., and Morrisey, E.E. (2018). Regeneration of the lung alveolus by an evolutionarily conserved epithelial progenitor. Nature 555, 251–255.

